# An ultrasensitive GRAB sensor for detecting extracellular ATP *in vitro* and *in vivo*

**DOI:** 10.1101/2021.02.24.432680

**Authors:** Zhaofa Wu, Kaikai He, Yue Chen, Hongyu Li, Sunlei Pan, Bohan Li, Tingting Liu, Huan Wang, Jiulin Du, Miao Jing, Yulong Li

**Author notes:** Correspondence (Z.W.); (Y.L.).

## Abstract

The purinergic transmitter ATP (adenosine 5’-triphosphate) plays an essential role in both the central and peripheral nervous systems, and the ability to directly measure extracellular ATP in real time will increase our understanding of its physiological functions. We developed an ultrasensitive <u>G</u>PC<u>R</u> <u>A</u>ctivation‒<u>B</u>ased ATP sensor called GRAB_ATP1.0_, with a robust fluorescence response to extracellular ATP when expressed in several cell types. This sensor has sub-second kinetics, ATP affinity in the range of tens of nanomolar, and can be used to localize ATP release with subcellular resolution. Using this sensor, we monitored ATP release under a variety of *in vitro* and *in vivo* conditions, including primary hippocampal neurons, a zebrafish model of injury-induced ATP release, and LPS-induced ATP-release events in individual astrocytes in the mouse cortex measured using *in vivo* two-photon imaging. Thus, the GRAB_ATP1.0_ sensor is a sensitive, versatile tool for monitoring ATP release and dynamics under both physiological and pathophysiological conditions.

## INTRODUCTION

Adenosine 5′-triphosphate (ATP) is a universal energy-storing molecule used by virtually all living organisms. In addition to its metabolic function intracellularly, growing evidence suggest that released ATP into the extracellular space can serve as a signaling molecule (termed purinergic transmitter) (Burnstock, 1972), by binding and activating ionotropic P2X receptors and metabotropic P2Y receptors (Abbracchio et al., 2006; Khakh and North, 2012). In the nervous system, a wide range of functions are regulated by ATP, including pain sensation (Burnstock, 1996; Collier et al., 1966), mechanosensory and chemosensory transduction (Burnstock, 2009; Gourine et al., 2005), and synaptic transmission (Burnstock, 2006). Notably, noxious stimuli in the central nervous system (e.g., injury, low osmolality, and inflammation) can trigger a sustained increase in extracellular ATP (Davalos et al., 2005; Wang et al., 2004), which is considered as a multi-target “danger” signal (Rodrigues et al., 2015). Not surprisingly, impaired ATP signaling has been associated with pathological processes (Burnstock, 2007, 2008; Cheffer et al., 2018). Despite the central role that ATP plays in both health and disease, the detailed mechanisms underlying the release and extracellular distribution of ATP are poorly understood, especially *in vivo*.

A significant number of advances in the last few decades culminated in a variety of techniques and tools for measuring extracellular ATP (Dale, 2021; Wu and Li, 2020). Unfortunately, despite their advantages, these techniques have several key limitations. For example, methods such as microdialysis, electrochemistry-based probes, reporter cells, and bioluminescent assays can measure ATP both *in vitro* and *in vivo* (Pellegatti et al., 2008), but are severely limited with respect to precisely detecting ATP due to their relatively low spatial and/or temporal resolution. On the other hand, fluorescent sensor‒based imaging can provide excellent spatiotemporal resolution (Giepmans et al., 2006), and several fluorescent protein‒based sensors have been developed for measuring extracellular ATP, including the recent ecAT3.10 (Conley et al., 2017) and pm-iATPSnFR (Lobas et al., 2019) sensors; however, these sensors are not compatible with measuring extracellular ATP *in vivo*, mainly due to their limited sensitivity and/or signal-to-noise ratio. A recently developed ATP sensor known as ATPOS (ATP Optical Sensor) has a high affinity for ATP and has been used to image extracellular ATP in the mouse cortex (Kitajima et al., 2020); however, this sensor must be injected as a recombinant protein (Kitajima et al., 2020), which can cause tissue damage and is difficult to measure ATP in a cell type specific manner. In addition to adapting soluble bacterial F_0_F_1_-ATP synthase as an ATP-binding protein (e.g., ecAT3.10, pm-iATPSnFR and ATPOS), the natural-evolved extracellular ATP “detectors”—ATP receptors—were also used to engineer ATP sensors. For example, taking advantage of the permeability to Ca^2+^ ions during ATP-gated P2X channel opening, versatile tools were developed by fusing the genetically-encoded Ca^2+^ indicators to the C-terminal of P2X subunits (Ollivier et al., 2021; Richler et al., 2008). These sensors display fast kinetics and/or sensitivity allowing to detect ATP release; however, it might be difficult to exclude that in some conditions, especially under *in vivo* systems, an ATP-P2X-independent activation of GCaMP6s may occur. Overall, the lack of genetically encoded tools that can sense a change in extracellular ATP concentration with high spatiotemporal resolution, high specificity, and high sensitivity has limited our ability to study purinergic signaling under both physiological and pathophysiological conditions.

Recently, our group and others developed a series of genetically encoded GPCR activation‒based (GRAB) sensors to measure a variety of neuromodulators—including acetylcholine (Jing et al., 2020; Jing et al., 2018), dopamine (Patriarchi et al., 2018; Patriarchi et al., 2020; Sun et al., 2018; Sun et al., 2020), norepinephrine (Feng et al., 2019), serotonin (Wan et al., 2020), and adenosine (Peng et al., 2020)—with high sensitivity, selectivity, and spatiotemporal resolution, providing the ability to monitor these neuromodulators in targeted cells under *in vivo* setting. Here, we report the development and application of a new GFP-based GRAB_ATP_ sensor using a P2Y receptor as the ATP-binding scaffold. This sensor, which we call GRAB_ATP1.0_ (short as ATP1.0), can be expressed in a wide range of cell types, producing a robust fluorescence response (with a ΔF/F_0_ of 500-1000%), and with high selectivity for both ATP and ADP; moreover, this sensor can be used to detect changes of extracellular ATP both *in vitro* and *in vivo* under a variety of conditions.

## RESULTS

### Development and characterization of a new GRAB sensor for detecting ATP

To develop a genetically encoded GRAB sensor for detecting ATP, we first systematically screened a series of candidate G protein‒couple receptors (GPCRs) known to be activated by ATP, including the human P2Y_1_, P2Y_2_, P2Y_4_, P2Y_11_, P2Y_12_, and P2Y_13_ receptors (Xing et al., 2016). Using these GPCRs as the scaffold, we inserted cpEGFP into the receptor flanked by short linker peptides at both the N- and C-terminus (Figure S1A); we selected the hP2Y_1_-based chimera ATP0.1 for further optimization based on its good membrane trafficking and high fluorescence response upon application of 100 μM ATP (Figure S1B). We then optimized the length and amino acid composition of the linkers between the hP2Y_1_ receptor and the cpEGFP moiety (Figure 1A) and identified the candidate with the largest fluorescence response (Figure 1B); we call this sensor GRAB_ATP1.0_ (hereafter referred to as ATP1.0). When expressed in HEK293T cells, ATP1.0 trafficked to the plasma membrane (Figure 1C) and produced a peak ΔF/F_0_ value of 500% in response to 100 μM extracellular ATP (Figure 1B and 1C). As a negative control, we also generated a mutant version of this sensor called ATP1.0mut, which contains the N283A mutation in the receptor’s ATP-binding pocket (Zhang et al., 2015), thus is non-sensitive to ATP (Figures S2 and S3).

**Figure 1.**
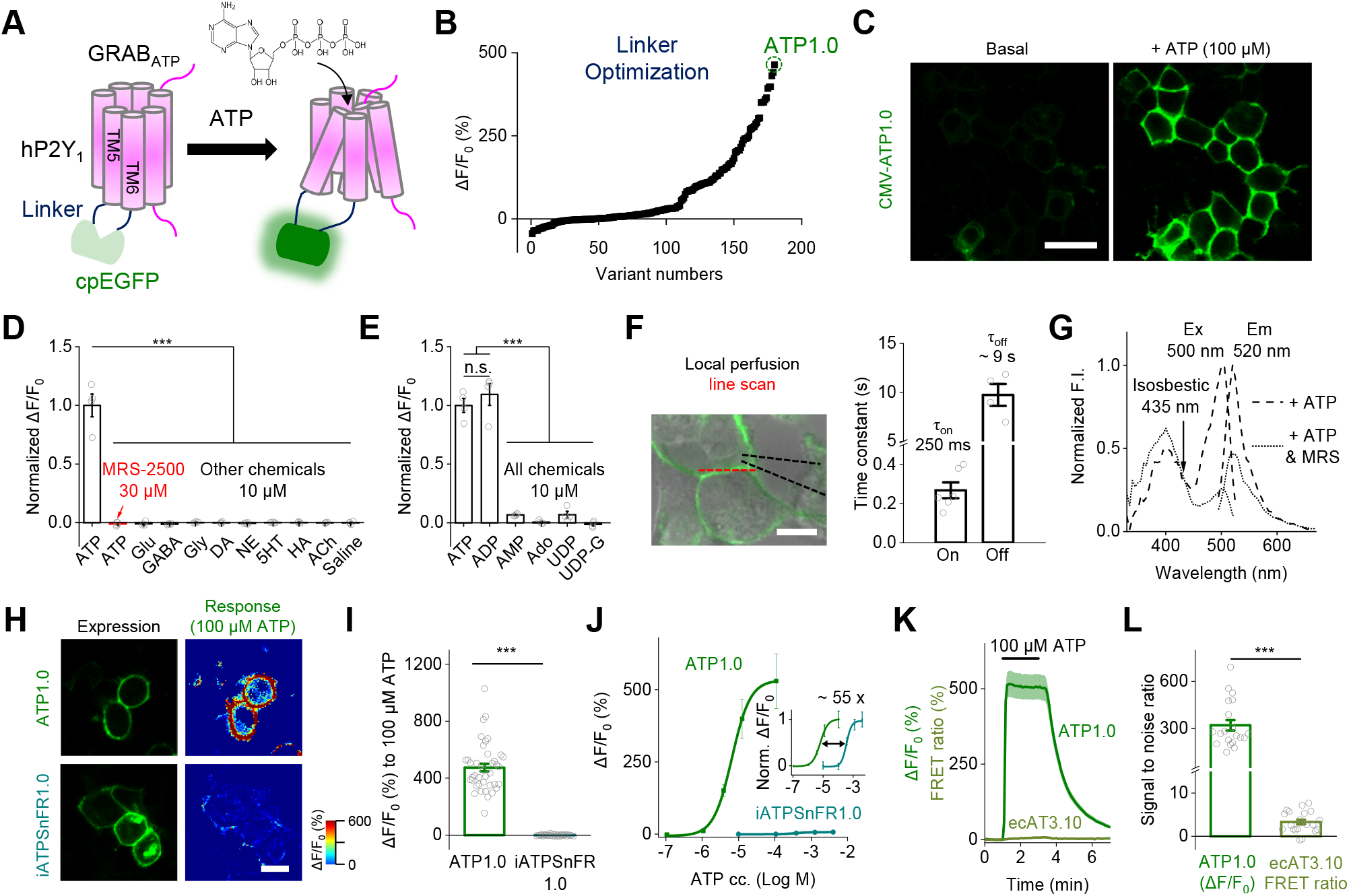
Design, optimization, and characterization of a genetically encoded GRABATP sensor. **(A)** Schematic drawing depicting the principle of GRAB-based ATP sensors designed using the human P2Y_1_ receptor as the scaffold coupled to circularly permuted enhanced GFP (cpEGFP). Binding of ATP induces a conformational change that increases the fluorescence signal. **(B)** Optimization of the N- and C-terminal linkers connecting the P2Y_1_ receptor and the cpEGFP moiety, yielding increasingly responsive ATP sensors. The sensor with the highest response to 100 μM ATP, GRAB_ATP1.0_ (ATP1.0), is indicated. **(C)** Example fluorescence images of HEK293T cells expressing the ATP1.0 sensor under basal conditions and in the presence of 100 μM ATP. **(D and E)** Summary of ΔF/F_0_ measured in ATP1.0-expressing HEK293T cells in the presence of the indicated compounds (each at 10 μM, except for MRS-2500, which was applied at 30 μM), normalized to the peak response measured in ATP; n = 4 independent wells each. ATP, adenosine triphosphate; MRS, MRS-2500; Glu, glutamate; GABA, γ-aminobutyric acid; Gly, glycine; DA, dopamine; NE, norepinephrine; 5-HT, 5-hydroxytryptamine (serotonin); HA, histamine; ACh, acetylcholine; ADP, adenosine diphosphate; AMP, adenosine monophosphate; Ado, adenosine; UDP, uridine diphosphate; UDP-G, UDP-glucose; Glu, glutamate. **(F)** Summary of the response kinetics of ATP1.0. Left, the experimental system in which ATP was locally puffed on an HEK293T cell expressing ATP1.0; a line-scan was used to measure the fluorescence response. Right, on kinetics was measured using a local puff of 100 μM ATP, and off kinetics was measured by a local puff of the P2Y_1_ receptor antagonist MRS-2500 in the presence of 10 μM ATP; n = 6 and 4 cells each for τ_on_ and τ_off_, respectively. **(G)** Excitation (Ex) and emission (Em) spectra of the ATP1.0 sensor in the presence of ATP (100 μM) or ATP (100 μM) together with MRS-2500 (300 μM). The isosbestic point at 435 nm is indicated. **(H and I)** GFP fluorescence images (left column) and pseudocolor images of the response (right column) measured in HEK293T cells expressing ATP1.0 (top row) or iATPSnFR1.0 (bottom row). Panel **(I)**shows the summary of the response to 100 μM ATP; n = 40 and 30 cells each for ATP1.0 and iATPSnFR1.0, respectively. **(J)** The peak fluorescence response measured in HEK293T cells expressing ATP1.0 or iATPSnFR1.0 plotted against the indicated concentrations of ATP; n = 10 and 20 cells each, respectively. Inset: the same data, normalized and re-plotted. **(K and L)** The fluorescence response (**K**) and signal-to-noise ratio (**L**) measured in HEK293T cells expressing ATP1.0 or the FRET-based ecAT3.10 sensor; where indicated, 100 μM ATP was applied; n = 20 cells each. The signal-to-noise ratio is defined as the peak response divided by the standard deviation prior to the application of ATP application. The scale bars represent 30 μm. The data in **(D)** and **(E)** were analyzed using a one-way ANOVA followed by Dunnett’s post-hoc test; the data in **(I)** and **(L)** were analyzed using the Student’s *t*-test. In this and subsequent figures, summary data are presented as the mean ± SEM; \*\*\**p*<0.001; n.s., not significant (*p*>0.05).

We then characterized the specificity, kinetics, brightness and spectrum of the ATP1.0 sensor. With respect to specificity, the ATP-induced response was fully blocked by the P2Y_1_ receptor antagonist MRS-2500, and no measurable response was produced by any other neurotransmitters or neuromodulators tested, including glutamate, GABA, glycine, dopamine, norepinephrine, serotonin, histamine, and acetylcholine (Figure 1D). ADP and ATP produced a similar response, whereas structurally similar purinergic molecules or derivatives such as AMP, adenosine, UDP, and UDP-glucose virtually produced no response (Figure 1E). ATP1.0 has rapid response kinetics, with a rise time constant (τ_on_) of ~250 milliseconds and a decay time constant (τ_off_) of ~9 seconds upon application of ATP and subsequent application of MRS-2500, respectively (Figure 1F). With respect to the sensor’s brightness, ATP increased the brightness of ATP1.0 to approximately 64% of the brightness measured in cells expressing an hP2Y_1_-EGFP fusion protein (Figure S4A). Finally, ATP1.0 shows similar spectrum as EGFP under one-photon excitation, with the excitation peak at ~500nm and emission peak at ~520nm (Figure. 1G).

To compare the performance of ATP1.0 with other extracellular ATP sensors, including single wavelength‒based iATPSnFR sensors (Lobas et al., 2019) and FRET-based ecAT3.10 sensors (Conley et al., 2017), we expressed these sensors in HEK293T cells and performed confocal imaging. Although ATP1.0 and iATPSnFR1.0 were expressed at similar levels at the plasma membrane expression (Figure 1H) and have similar brightness (data not shown), cells expressing ATP1.0 had a 50-fold larger dynamic range to ATP compared to cells expressing iATPSnFR1.0 (Figure 1H-1J). Moreover, compared to cells expressing ATP1.0, cells expressing ecAT3.10 had an extremely small response (Figure 1K) and a significantly smaller signal-to-noise ratio (Figure 1L).

Next, we examined the performance of ATP1.0 in cultured rat primary astrocytes using an adeno-associated virus (AAV) expressing the sensor under the control of the astrocyte-specific GfaABC1D promoter (Lee et al., 2008). We found that ATP1.0 was widely distributed throughout the plasma membrane, including the soma and cell processes (Figures 2A and S5A). Similarly, when expressed in cultured cortical neurons under the control of the neuron-specific hSyn promotor, ATP1.0 was widely distributed throughout the plasma membrane, including the soma and neurites (Figures 2D and S5B). Both astrocytic and neuronally expressed ATP1.0 responded robustly to ATP application, with peak ΔF/F_0_ values of approximately 1000% and 780%, respectively (Figure 2A-2F). Moreover, the ATP-induced fluorescence response was blocked by the P2Y_1_ receptor antagonist MRS-2500 (Figure 2G and 2H), and virtually no response was observed in neurons expressing the control ATP1.0mut sensor (Figure S2B). In addition, similar to our results obtained with HEK293T cells, ATP1.0 expressed in neurons responded to both ATP and ADP, but did not respond to AMP, adenosine, UTP, or GTP (Figure 2G-2I). Importantly, the ATP1.0 sensor was stable at the cell surface, as we observed no decrease in fluorescence in ATP1.0-expressing neurons during a 2-hour application of 10 μM ATP (Figure 2J-2L).

**Figure 2.**
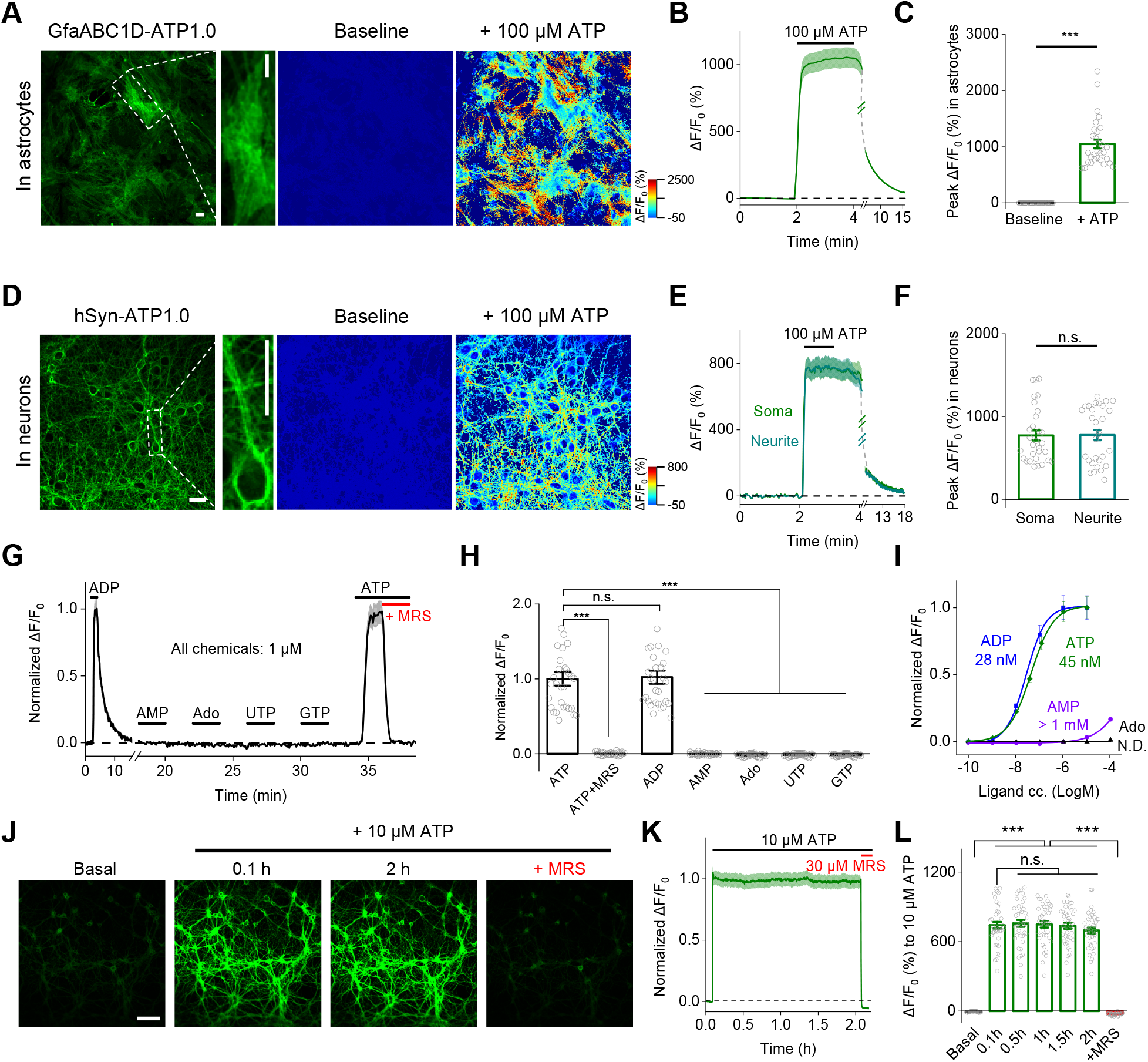
Characterization of the ATP1.0 sensor in primary cultured astrocytes and neurons. **(A-C)** ATP1.0 was expressed in cultured cortical astrocytes and measured using confocal imaging. **(A)** Raw GFP fluorescence image (left) and pseudocolor images of the baseline and peak response (ΔF/F_0_) to 100 μM ATP. **(B)** Time course of ΔF/F_0_; 100 μM ATP was applied where indicated. **(C)** Summary of the peak ΔF/F_0_ measured before and after application of 100 μM ATP; n = 30 ROIs each from 3 coverslips. **(D-F)** Same as **(A-C)**, except ATP1.0 was expressed in cultured rat cortical neurons; n = 30 ROIs each from 3 coverslips. **(G-I)** Normalized ΔF/F_0_ measured in cultured neurons expressing ATP1.0, showing an example trace **(G)**, summary data **(H)**, and dose-response curves with corresponding EC_50_ values **(I)**. UTP, uridine triphosphate; GTP, guanosine triphosphate; N.D., not determine; n = 30 ROIs from 3 coverslips **(H)**. **(J-L)** Fluorescence image **(J)**, trace **(J)**, and summary **(L)** of ATP1.0 expressed in cultured hippocampal neurons during a 2-hour application of 10 μM ATP; n = 40 neurons from 2 coverslips. Scale bars represent 30 μm **(A and D)**and 100 μm **(J)**. The data in **(C)** and **(F)** were analyzed using the Student’s *t*-test; the data in **(H)** and **(L)**were analyzed using a one-way ANOVA followed by Dunnett’s post-hoc test. \*\*\**p*<0.001; n.s., not significant.

Taken together, these results indicate that the ATP1.0 sensor is suitable for the use in several cell types, providing a sensitive, specific, and stable fluorescence increase in response to extracellular ATP.

### ATP1.0 can be used to monitor the release of ATP from cultured hippocampal neurons

Next, we examined whether the ATP1.0 sensor could be used to detect the release of endogenous ATP in neuron-glia co-cultures (Figure 3A), a widely used system for studying ATP signaling (Fields, 2011; Koizumi et al., 2003; Zhang et al., 2003). First, we tested whether ATP1.0 could detect stimulus-evoked ATP release. In the brain, ATP is released in response to mechanical stimulation and cell swelling (Newman, 2001; Xia et al., 2012). To induce a mechanical stimulus, we pressed a glass pipette against the cultured cells; when the ATP1.0 signal increased, we then removed the pipette to end the stimulus. We found that mechanical stimulation induced a rapid, localized increase in ΔF/F_0_, reflecting the release of ATP (Figure 3B). To induce cell swelling, we bathed the cells in a hypotonic solution (130 mOsm/kg); within one minute, a robust increase in ΔF/F_0_ was observed (Figure 3D). Importantly, the responses induced by both stimuli were abolished by the application of MRS-2500 and were absent in cells expressing the control ATP1.0mut sensor (Figure 3B-3E), confirming the specificity of ATP1.0. We also found that the hypotonic stimulus‒induced release of ATP may not require classical SNARE-dependent vesicular releasing machinery, as expressing tetanus toxin light chain (TeNT), which cleaves synaptobrevin and prevents exocytosis (Patterson et al., 2010; Schiavo et al., 1992), had no effect on the response in cells expressing hSyn-ATP1.0 (Figure 3F1 and 3G1); as a control, expressing TeNT abolished the stimulation-evoked release of glutamate (Glu) release measured using the Glu sensor SF-iGluSnFR.A184V (Figure 3F2 and 3G2).

**Figure 3.**
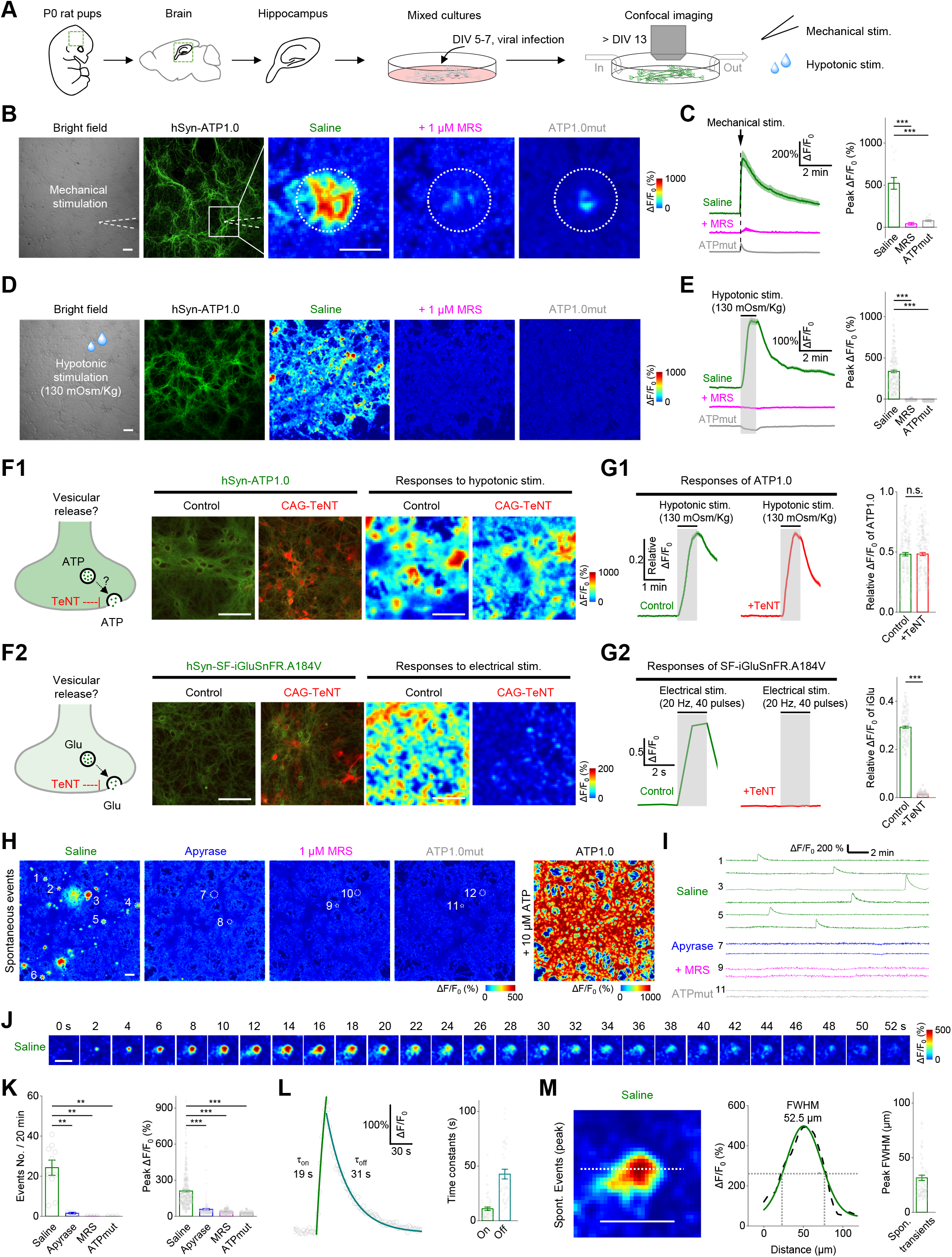
Release of endogenous ATP in primary hippocampal cultures. **(A)** Schematic diagram depicting the experimental protocol in which primary hippocampal neurons are cultured and infected with an AAV encoding ATP1.0 or ATP1.0mut under the control of the hSyn promoter, followed by confocal fluorescence microscopy during various stimuli. DIV, days *in vitro*. **(B-E)** Bright field images, GFP fluorescence images, and pseudocolor images **(B and D)**, and average traces **(C and E, left)** of the fluorescence response of ATP1.0 or ATP1.0mut measured in saline or 1 μM MRS-2500 (MRS). The white dashed circles in **(B)** indicate the 150-μm diameter ROI used for analysis, and the white dashed lines in **(B)** indicate the location of the electrode used for mechanical stimulation. The summary data **(C and E, right)** represent 13-20 ROIs from 3 coverslips **(C)** and 170-214 ROIs from 3-4 coverslips **(E)**. **(F1 and G1)** Fluorescence images of ATP1.0 (green) and EBFP2-iP2A-TeNT (red) **(F1)**, pseudocolor images **(F1)**, average traces **(G1, left)**, and summary data **(G1, right)**; n = 217-227 ROIs from 4 coverslips each. **(F2 and G2)** Fluorescence images of SF-iGluSnFR.A184V (green) and TeNT-BFP2 (red) **(F2)**, pseudocolor images **(F2)**, average traces **(G2, left panels)**, and summary data **(G2, right)**; n = 171 ROIs from 3 coverslips each. **(H)** Cumulative transient change in ATP1.0 or ATP1.0mut fluorescence measured during 20 min of recording in saline, apyrase (30 U/ml), or 1 μM MRS-2500. The white dashed circles indicate the ROIs used for the analyses in **(I)**. **(I)** Exemplar traces of ΔF/F_0_ measured under the indicated conditions. **(J)** Exemplar time-lapse pseudocolor images captured in saline. **(K)** Quantification of the number of events per 20 min (left) and the peak fluorescence response (right) in neurons expressing ATP1.0 or ATP1.0mut; n = 114-363 ROIs from 3-10 coverslips. **(L and M)** Kinetics profile **(L)** and spatial profile **(M)** of the change in ATP1.0 fluorescence measured in saline. The summary in **(L)** and **(M)** data represent 54 events and 128 events, respectively, from 4 coverslips. Scale bars represent 100 μm. The data in **(C)** and **(E)** were analyzed using a one-way ANOVA followed by Dunnett’s post-hoc test; the data in **(G)** were analyzed using the Student’s *t*-test. the data in **(K)** were analyzed using a one-way ANOVA with Bonferroni correction. \*\**p*<0.01; \*\*\**p*<0.001; n.s., not significant.

In addition to stimulus-evoked ATP release, we also observed spontaneous, localized, transient ATP1.0 signals in our neuron-glia co-cultures even in the absence of external stimulation (Figure 3H and 3I). In the 1.6-mm^2^ imaging field, these events occurred at a rate of 1.2/min and had an average peak ΔF/F_0_ of approximately 210% (Figure 3K). The average rise time (τ_on_) and decay time (τ_off_) of spontaneous ATP-releasing events were 11 s and 43 s (Fig. 3L), respectively. The average diameter of spontaneous ATP-releasing events was 32 μm based on our analysis of full width at half maximum (FWHM) (Fig. 3M). In contrast, no spontaneous events were observed in the presence of MRS-2500 or in cells expressing ATP1.0mut (Figure 3H, 3I and 3K). To confirm that the ATP1.0 signal reflects extracellular ATP dynamics, we imaged cells in the presence of the ATP degrading enzyme apyrase. We observed that apyrase (30 U/ml) treatment significantly blocked spontaneous events (Figure 3H, 3I and 3K).

### ATP1.0 can be used to measure the injury-induced *in vivo* propagation of ATP in zebrafish larvae

Having shown that the ATP1.0 sensor is suitable for use in *in vitro* systems, we then examined whether it could be applied to monitor ATP in *in vivo* systems such as zebrafish. We therefore transiently expressed either ATP1.0 or ATP1.0mut in neurons of larval zebrafish under the control of the neuron-specific *elval3* promoter (Figure 4A and 4B). Local puffing of ATP, but not saline, elicited a robust transient increase in ΔF/F_0_ in the optic tectum. These signals were blocked by MRS-2500 and not observed in zebrafish larvae expressing the control ATP1.0mut sensor (Figure 4C).

**Figure 4.**
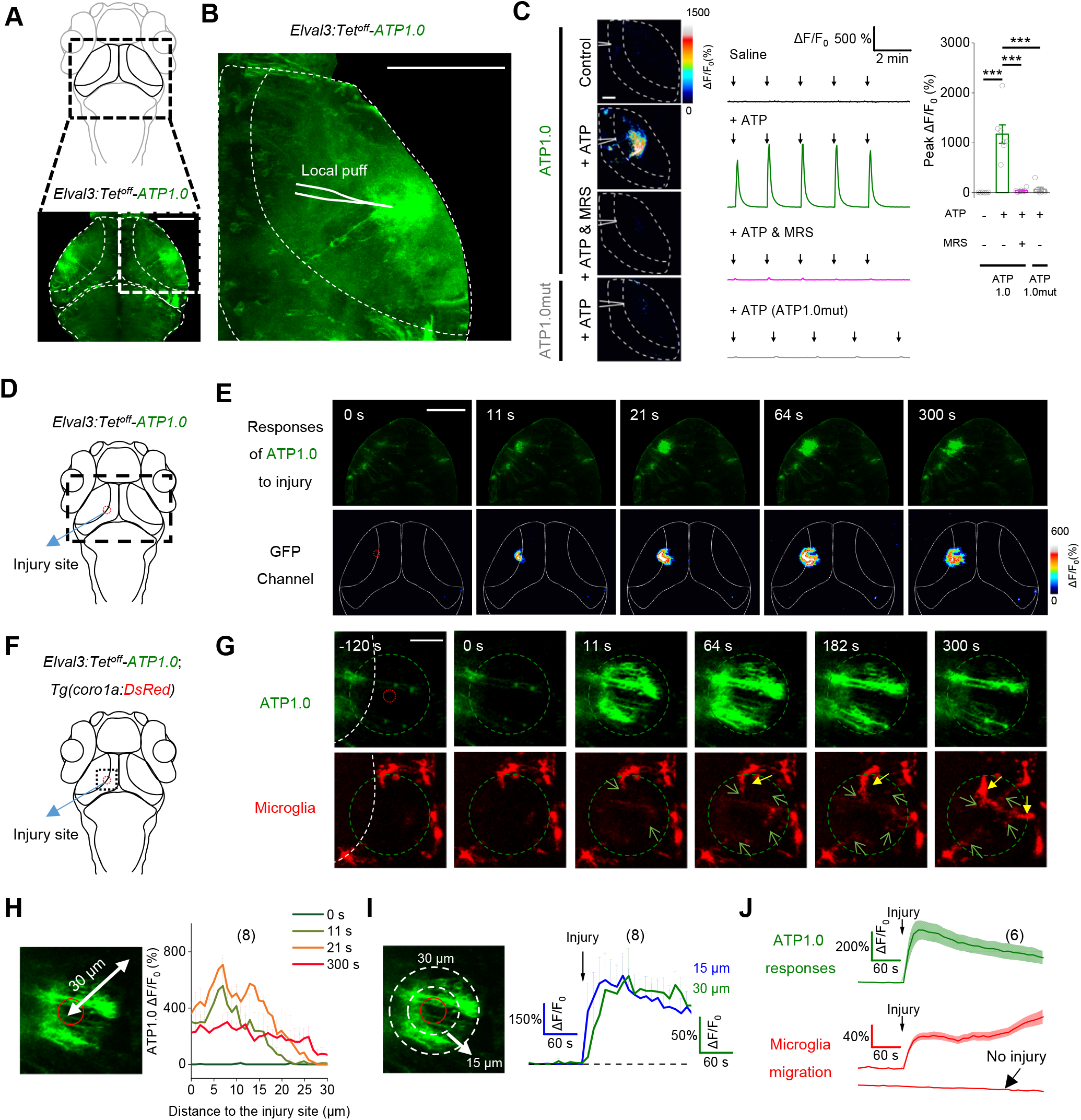
ATP1.0 reveals in vivo ATP release induced by injury in a zebrafish model. **(A and B)** Schematic diagram depicting *in vivo* confocal imaging of fluorescence changes induced by a localized puff (via a micropipette; see inset) of various compounds in the optic tectum of zebrafish larvae expressing ATP1.0 (*Elval3: Tet^off^-ATP1.0*) or ATP1.0mut (*Elval3: Tet^off^-ATP1.0mut*). **(C)** Example fluorescence images (left), traces (middle), and summary (right) showing the response of ATP1.0 or ATP1.0mut to the indicated compounds. Arrows indicate the localized application of saline (Control) or ATP (5 mM). Where indicated, MRS-2500 (90 μM) was applied; n = 6-7 fishes. **(D)** Schematic diagram depicting confocal imaging of ATP1.0 responses before and after two-photon laser ablation (i.e., injury) in the optic tectum of zebrafish larvae expressing ATP1.0. The red dashed circle indicates the region of laser ablation, and the black dashed rectangle indicates the imaging region shown in **(E)**. **(E)** Time-lapse pseudocolor images showing the response of ATP1.0 to laser ablation in the optic tectum. The laser ablation was performed at time 0 s and lasted for 7-sec, and ATP1.0 fluorescence was imaged beginning 2 min before laser ablation. **(F)** Schematic diagram showing dual-color confocal imaging of ATP release and microglial migration before and after laser ablation in transgenic zebrafish *Tg(coro1a: DsRed)* larvae expressing ATP1.0. In *Tg(coro1a: DsRed)* larvae, the microglia expresses DsRed. The red dashed circle indicates the region of laser ablation, and the black dashed rectangle indicates imaging region. **(G)** *In vivo* time-lapse confocal images showing the migration of microglia (red) and the change in ATP1.0 fluorescence (green) before and after laser ablation (start at time 0 s). The green dashed circle indicates the boundary of the ATP wave at 300 s, and the signal measured in the green dashed circle was used for the analysis in **(J)**. Green arrows indicate the protrusions of microglia; solid yellow arrows indicate the cell bodies of microglia. **(H)** Summary of the distance between the ATP1.0 response and the site of injury measured at 0, 11, 21, and 300 s after injury. **(I)** Time course of the ATP1.0 response measured 15 and 30 μm from the site of laser ablation. The arrow indicates the beginning of the 7-sec laser ablation. **(J)** Time course of the ATP1.0 response (green) and the microglia migration (red) before and after laser ablation (vertical arrow); also shown is a trace of DsRed fluorescence measured in the absence of laser ablation. 30-μm diameter ROIs were used for analysis. Scale bars represent 40 μm **(B)** and **(C)**, 100 μm **(E)**, and 30 μm **(G)**. The numbers in parentheses in **(H-J)** represent the number of zebrafish larvae in each group. \*\*\**p*<0.001 (Student’s *t-*test).

Next, we examined whether ATP1.0 could be used to measure the release of endogenous ATP in live zebrafish. It is known that ATP signaling plays key roles in promoting the migration of microglia to injury site (Li et al., 2012; Sieger et al., 2012). We found that injury induced by laser ablation in the optic tectum caused a robust increase in fluorescence in ATP1.0-expressing zebrafish (Figure 4D and 4E). Moreover, the response propagated in a radial pattern outward from the site of injury (Figure 4E, 4H, and 4I). Next, we simultaneously monitored ATP release and the migration of microglia by expressing ATP1.0 in the optic tectum of a transgenic zebrafish line in which the microglia are labeled with the red fluorescent protein DsRed (Figure 4F). We found that following laser ablation, microglia gradually migrated to the site of injury along the path of ATP propagation measured using ATP1.0 (Figure 4G and 4J). Thus, our ATP1.0 sensor is well-suited for *in vivo* application in zebrafish larvae, providing high spatiotemporal resolution.

### ATP1.0 can be used to monitor localized ATP release during LPS-induced systemic inflammation in mice

Purinergic signaling molecules, including ATP, are considered critical extracellular messengers in response to acute and chronic inflammation, acting via paracrine or autocrine processes on immune cells in the peripheral nervous system and on neurons and glia cells in the central nervous system (Idzko et al., 2014). To date, however, the pattern by which ATP is released during systemic inflammation, as well as the relationship between this release and inflammatory status, are poorly understood. We therefore used a mouse model of systemic inflammation induced by an intraperitoneal injection of bacterial lipopolysaccharides (LPS; 10 mg/kg), and directly observed ATP dynamics in the visual cortex using two-photon imaging of ATP1.0 fluorescence (Figure 5A); this inflammation model caused a robust increase in expression of the inflammatory cytokines IL-1β and IL-10 in the brain (Figure S6). Twenty-four hours after LPS injection, we observed multiple localized ATP-release events in the cortex, with a frequency of approximately 5-10 events/min measured during 20 minutes of recording (Figure 5B2 and 5D). In contrast, fewer events occurred prior to LPS injection (data not shown), in saline-injected controls (Figure 5B1, 5C and 5E), and no events were observed in LPS-injected mice expressing the mutant ATP1.0mut sensor (Figure 5B3 and 5C).

**Figure 5.**
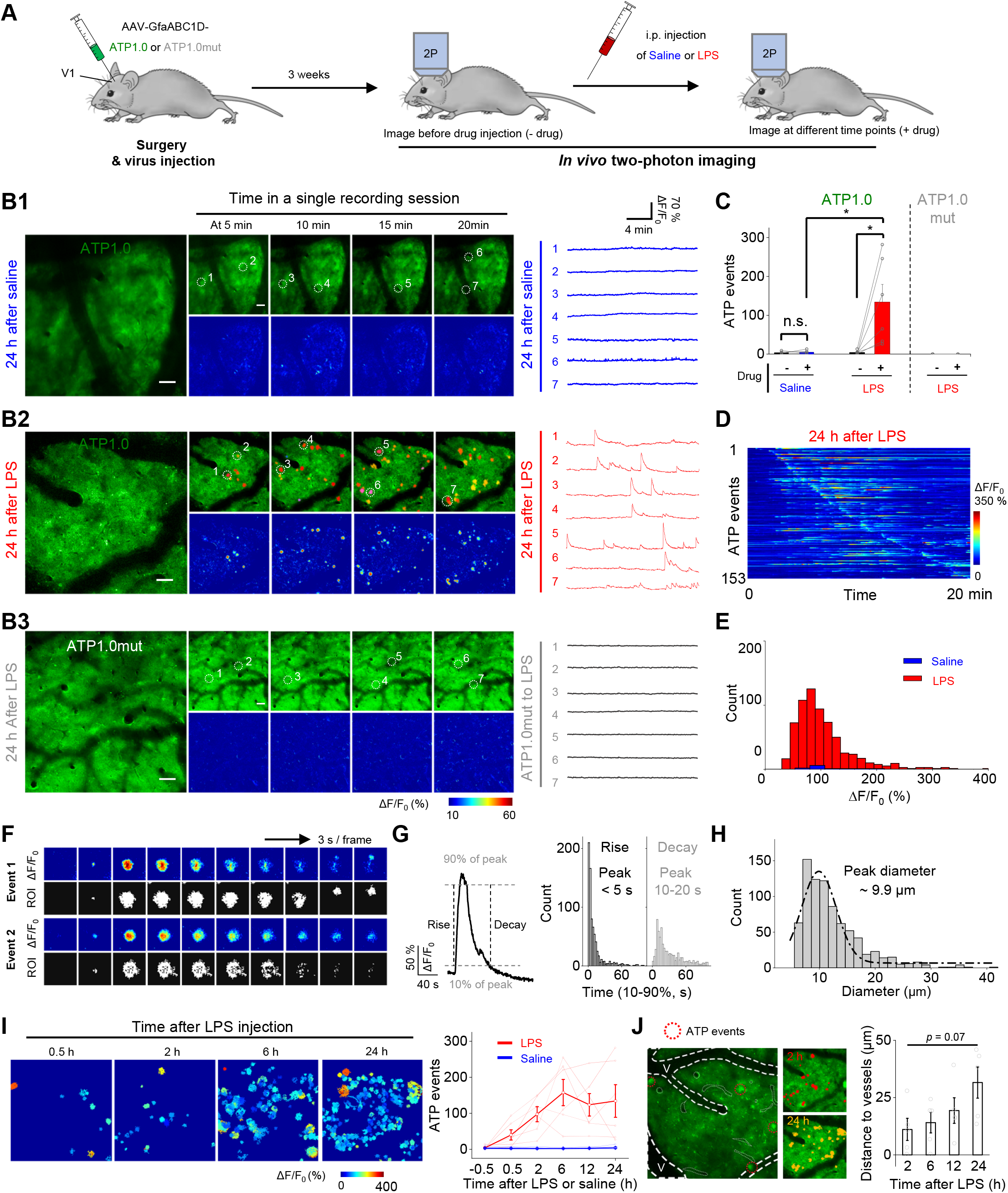
ATP1.0 reveals localized ATP-release events measured in the mouse brain following systemic inflammation induced by an injection of LPS. **(A)** Schematic diagram depicting the experimental protocol in which an AAV encoding either ATP1.0 or ATP1.0mut under the control of the GfaABC1D promoter is injected into the mouse visual cortex (V1), followed by two-photon imaging through a cranial window at various times after an intraperitoneal (i.p.) injection of saline or lipopolysaccharides (LPS, 10 mg/kg). **(B1 and B2)** Exemplar fluorescence images, pseudocolor images, and individual traces of the fluorescence response of ATP1.0 measured 24 h after saline **(B1)** or LPS **(B2)** injection using the indicated regions of interest (white dashed circles) identified using AQuA software overlay. **(B3)** Same as **(B2)**, except the ATP1.0mut sensor is expressed. **(C)** Summary of the number of localized ATP events measured during a 20-min recording before (−) and after (+) saline or LPS injection; n = 5-6 mice each. **(D)** Pseudocolor images showing all the identified ATP events in an exemplar ATP1.0-expressing mice after 24 h LPS injection. The number of identified ATP events from one mouse is shown on the y-axis. **(E)** Distribution of the peak fluorescence response (ΔF/F_0_) of the localized ATP events measured in ATP1.0-expressing mice 24 h after LPS (red) or saline (blue) injection; n = 805 events from 6 mice and 25 events from 5 mice, respectively. **(F)** Detailed analysis of the properties of two individual localized ATP events shown as pseudocolor images of ΔF/F_0_ and the corresponding ROIs identified using AQuA software at 3-s intervals. **(G)** Left, a representative trace (averaged from 50 peak-aligned events) showing the rise and decay kinetics of the event, defined the time between 10% and 90% of the baseline to peak. Right, summary of rise and decay times; n = 805 events from 6 mice. **(H)** Distribution of the size of the individual events measured in ATP1.0-expressing mice 24 h after LPS injection; n = 805 events from 6 mice. **(I)** Left, representative images showing ATP-release events (indicated as ROIs) in ATP1.0-expressing mice at the indicated times after LPS injection. Right, summary of the number of ATP-release events measured during 20-min recordings at the indicated times after LPS or saline injection. The data from each individual mouse and the average data are shown; n = 6 and 5 mice for the LPS and saline groups, respectively. **(J)** Left, representative image showing early ATP-release events red dashed circles) located near the blood vessel (V, indicated by white dashed lines). Middle, images taken 2 hours (red) and 24 hours (yellow) after LPS injection. Right, summary of the distance between the events and the blood vessel at the indicated times after LPS injection; n = 6 mice. Scale bars represent 50 μm **(B)**. The data in **(C)** were analyzed using the Student’s *t*-test; the data in **(J)** were analyzed using a one-way ANOVA followed by Dunnett’s post-hoc test. \**p*<0.05; n.s., not significant.

Next, we used the Astrocyte Quantitative Analysis (AQuA) software (Wang et al., 2019) to characterize the individual events. The ATP-release events had broadly distributed signal kinetics, although the majority of events have a relatively fast rise time (<5 s) and a slower decay time (10-20 s) (Figure 5F and 5G). In addition, the events had a spatially selective pattern, with an average signal diameter (determined using the maximum diameter of each event) of approximately 9.9 μm (Figure 5H), smaller than the average diameter of a typical astrocyte (10-20 μm) (Chai et al., 2017). To detailly examine the correlation between the ATP-release events and the progression of inflammation, we recorded cortical ATP events at various time points after LPS injection. We found an increase in ATP-release events within 30 min of LPS injection, and the number of events increased progressively with time, reaching a plateau 6 hours after injection; in contrast, no events were detected in saline-injected mice at any time point up to 24 hours (Figure 5I). Interestingly, an analysis of the location of the ATP-release events within the cortex revealed that the early events occurred relatively close to the blood vessels, and the distance between the events and the nearest vessels increased with time (Figure 5J). These data suggest that the brain can sense inflammation and respond in the form of spatially selective ATP-release events, demonstrating that the ATP1.0 sensor is compatible with *in vivo* imaging in mice, with unprecedented sensitivity and spatiotemporal resolution.

## DISCUSSION

Here, we report the development and characterization of a new, ultrasensitive, genetically encoded ATP sensor called GRAB_ATP1.0_. We also show that this sensor can be expressed reliably in a variety of cell types, including cell lines, astrocytes, and neurons, providing a robust tool for measuring extracellular ATP. Moreover, we show that this sensor can be used to visualize the real-time release of endogenous ATP *in vitro*, as well as ATP signaling in two *in vivo* models under several conditions.

Our GRAB_ATP_ sensors have at least four distinct advantages over other sensors with respect to monitoring the dynamics of extracellular ATP. First, ATP1.0 has extremely high sensitivity for extracellular ATP compared to other ATP sensors such as the recently developed, genetically-encoded single-wavelength ATP sensor, iATPSnFR1.0. When expressed in HEK293T cells, GRAB_ATP1.0_ displayed an EC_50_ of ~6.7 μM with a maximum ΔF/F_0_ of ~500% (Figure. 1J). Under the same condition, iATPSnFR1.0 displayed excellent plasma membrane localization (Figure 1H), yielding an EC_50_ ~381 μM (Figure 1G), which was consistent with the published data (Lobas et al., 2019). However, the maximum ΔF/F_0_ of iATPSnFR1.0 is ~10%, ~10-fold lower than the reported data (Lobas et al., 2019), presumably because of different imaging conditions. Curiously, we found that the GRAB_ATP1.0_ exhibits apparently different affinities to ATP in HEK293T cells (apparent EC_50_ ~6.7 μM) vs. neurons (apparent EC_50_ ~45 nM). One reason we speculated is due to the existence of enzymes that degraded the ATP in cultured HEK293T cells, which reduced the apparent affinities. Given the high sensitivity of GRAB_ATP1.0_ sensors, particularly when expressed in neurons and astrocytes, ATP1.0 will be useful for studying both pathological and physiological processes. Second, the ATP1.0 sensor is genetically encoded and can be expressed selectively in a variety of cell types, providing cell type‒specific measurements of ATP transmission. Third, ATP1.0 has high spatial resolution, suitable for measuring highly localized, transient ATP-release events in hippocampal cultures and in the mouse cortex. Lastly, our results demonstrated ATP1.0 can be used to monitor ATP dynamics *in vivo* using a variety of animal models, including zebrafish and mice.

Despite these advantages of genetically encoded GRAB_ATP_ sensors, a potential caveat is that ATP1.0 is based on the scaffold P2Y_1_ receptor (Waldo et al., 2002) and therefore responds to both ATP and ADP. Given that ATP and ADP may regulate distinct processes, particularly in the peripheral nervous system (Gaarder et al., 1961), next-generation GRAB_ATP_ sensors should be developed with improved molecular specificity, for example by engineering the GPCR scaffold to increase the sensor’s selectivity for ATP over ADP, and vice versa. Alternatively, other P2Y receptors, such as P2Y_11_ and P2Y_12_, which are more specific for ATP (Communi et al., 1997) and ADP (Hollopeter et al., 2001), respectively, can be used as scaffolds in developing future ATP or ADP sensors (Fig. S1B).

In hippocampal cultures, the ATP1.0 sensor readily resolved both evoked and spontaneous ATP release. Moreover, our study revealed that the hypotonicity‒induced ATP release was not sensitive to TeNT, supporting a non‒ vesicular mechanism of ATP release (Lazarowski, 2012). Interestingly, several molecules are proposed to mediate stimulus-induced ATP release (Taruno, 2018), including calcium homeostasis modulator (CALHM) (Taruno et al., 2013), pannexin/connexin, P2X7 receptors (Pellegatti et al., 2005), Leucine Rich Repeat Containing 8 VRAC Subunit A (LRRC8A)/SWELL1 (Qiu et al., 2014; Voss et al., 2014) and SLCO2A1 (Sabirov et al., 2017). We anticipate the new developed ATP1.0 sensor will provide a good tool to further dissect the relative contributions of these channels on ATP release under different stimulation conditions.

By combining the ATP1.0 sensor with *in vivo* two-photon imaging, we detected highly localized ATP-release events in the mouse brain following a systemic injection of LPS, and we found that these events were smaller in size than the diameter of a single astrocyte (Chai et al., 2017), indicating that the brain can sense systemic inflammation and respond with ATP signaling at cellular level. Further combining ATP1.0 imaging with genetic and pharmacological tools may facilitate the identification of cell types and molecules required for ATP signaling during these processes. A growing body of experimental evidence suggests that neuroinflammation is a key pathological event triggering and perpetuating the neurodegenerative processes associated with many neurological diseases, including Alzheimer's disease, Parkinson's disease, and amyotrophic lateral sclerosis (Amor et al., 2014; Nguyen et al., 2002). Thus, our GRAB_ATP_ sensor can be a powerful tool for studying dynamic changes in ATP release and the role of these changes in the neuroinflammatory processes that underlie neurodegeneration.

ATP plays an important role in neuron-glia interactions, which has complex interaction with other signaling such as calcium or glutamate. For example, the release of ATP can trigger calcium waves in astrocytes and affect neuronal glutamate release (Bazargani and Attwell, 2016; Fields and Burnstock, 2006; Guthrie et al., 1999; Illes et al., 2019; Zhang et al., 2003). Thus, the ATP1.0 sensor can be combined with a spectrally compatible calcium indicator, glutamate sensor, and/or other fluorescent indicators, providing an orthogonal readout of ATP with extremely high spatial and temporal resolution, yielding new insights into the role of ATP signaling under both physiological and pathophysiological processes.

## ACKNOWLEDGMENTS

We thank the members of the Li lab for helpful suggestions and comments. This research was supported by the Beijing Municipal Science & Technology Commission (Z181100001318002 and Z181100001518004 to Y.L., and the Beijing Nova Program Z20111000680000 to M.J.); grants from the NSFC (81821092 and 92032000 to Y.L.); grants from National Key R&D Program of China (2020YFE0204000 to Y.L. and 2019YFA0801603 to J.D.); and grants from the Peking-Tsinghua Center for Life Sciences and the State Key Laboratory of Membrane Biology at Peking University School of Life Sciences (to Y.L.). Z.W. is supported by the Boehringer Ingelheim-Peking University Postdoctoral Program. We thank Xiaoguang Lei at PKU-CLS and the National Center for Protein Sciences at Peking University for their support and assistance using the Opera Phenix high-content screening system.

## AUTHOR CONTRIBUTIONS

Y.L. supervised the project. Z.W. and K.H. performed the experiments related to the development, optimization, and characterization of the sensors in cultured cells, with contributions from S.P., B.L., and H.W. Z.W. performed the imaging of ATP release in cultured cells. H.L. and T.L. performed the *in vivo* zebrafish experiments under the supervision of J.D. Y.C. and M.J. performed the *in vivo* two-photon imaging experiments in mice. All authors contributed to the interpretation and analysis of the data. Z.W. and Y.L. wrote the manuscript with input from all authors.

## SUPPLEMENTAL INFORMATION

**Figure S1.**
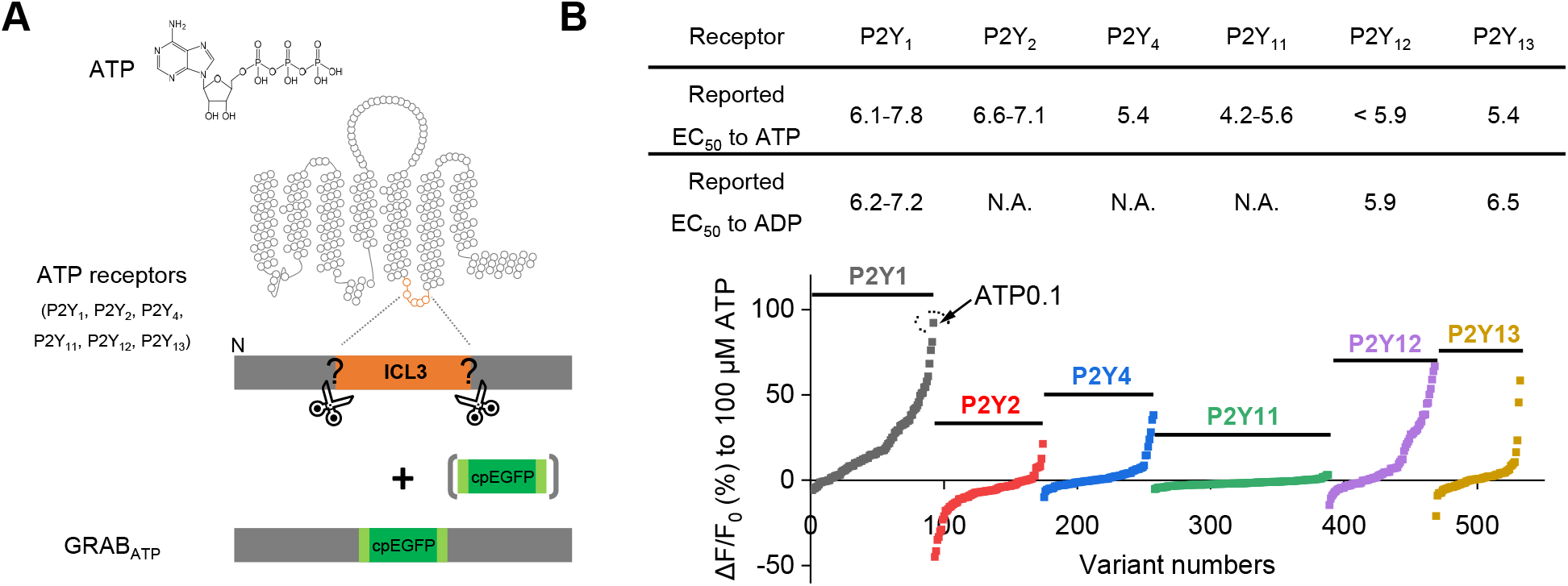
Selection of a GPCR scaffold for designing a genetically engineered GRAB-based ATP sensor. **(A)** Schematic diagram depicting the strategy for screening candidate GPCR scaffolds. **(B)** Upper panel, summary of the reported EC_50_ values of six human P2Y GPCRs for ATP and ADP (https://www.guidetopharmacology.org/); bottom panel, selection of the cpEGFP insertion site in the six candidates based on the fluorescence response of each variant to 100 μM ATP. The final sensor, ATP1.0, is based on the P2Y_1_ receptor and is indicated. ICL3, third intracellular loop; GPCR, G protein‒coupled receptor; N.A., not available.

**Figure S2.**
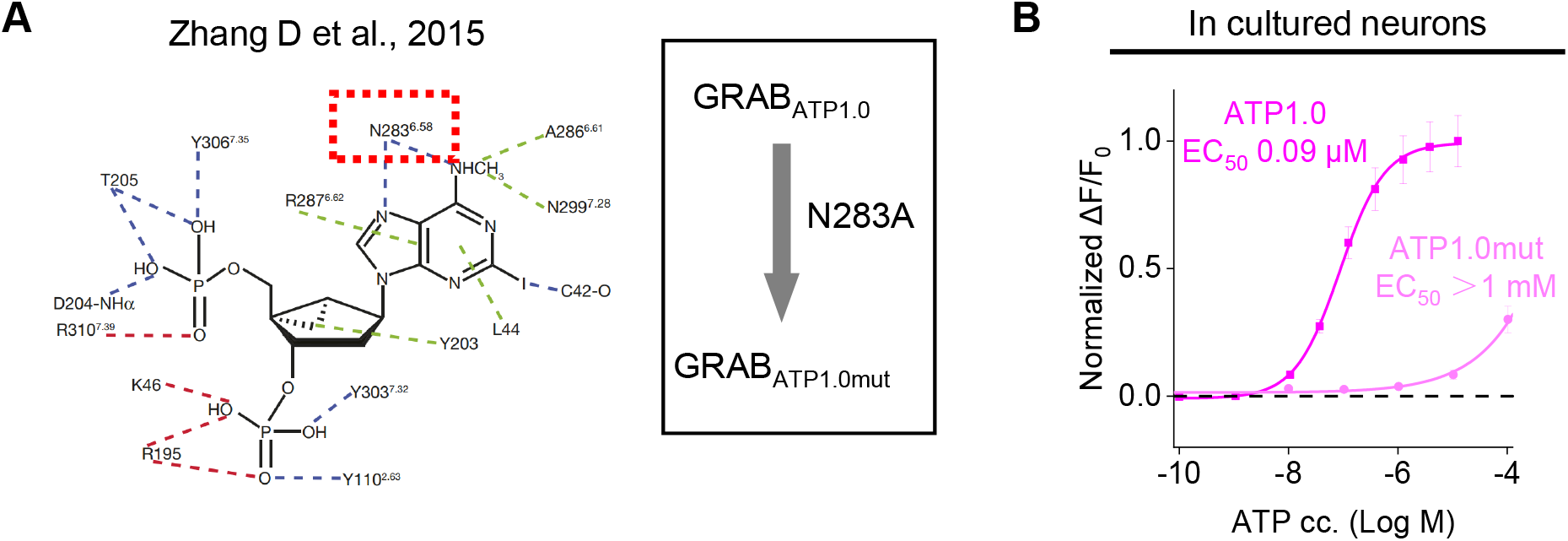
Design of an ATP-insensitive mutant sensor. **(A)** Location of N283, a key residue in the hP2Y_1_ receptor for ligand binding; this residue in ATP1.0 was mutated to an alanine (N283A mutation), resulting in the ATP1.0mut sensor. **(B)** Normalized dose-dependent fluorescence changes in neurons expressing either ATP1.0 or ATP1.0mut-expressing measured in response to ATP. Each point represents the average response measured in 12-14 ROIs.

**Figure S3.**
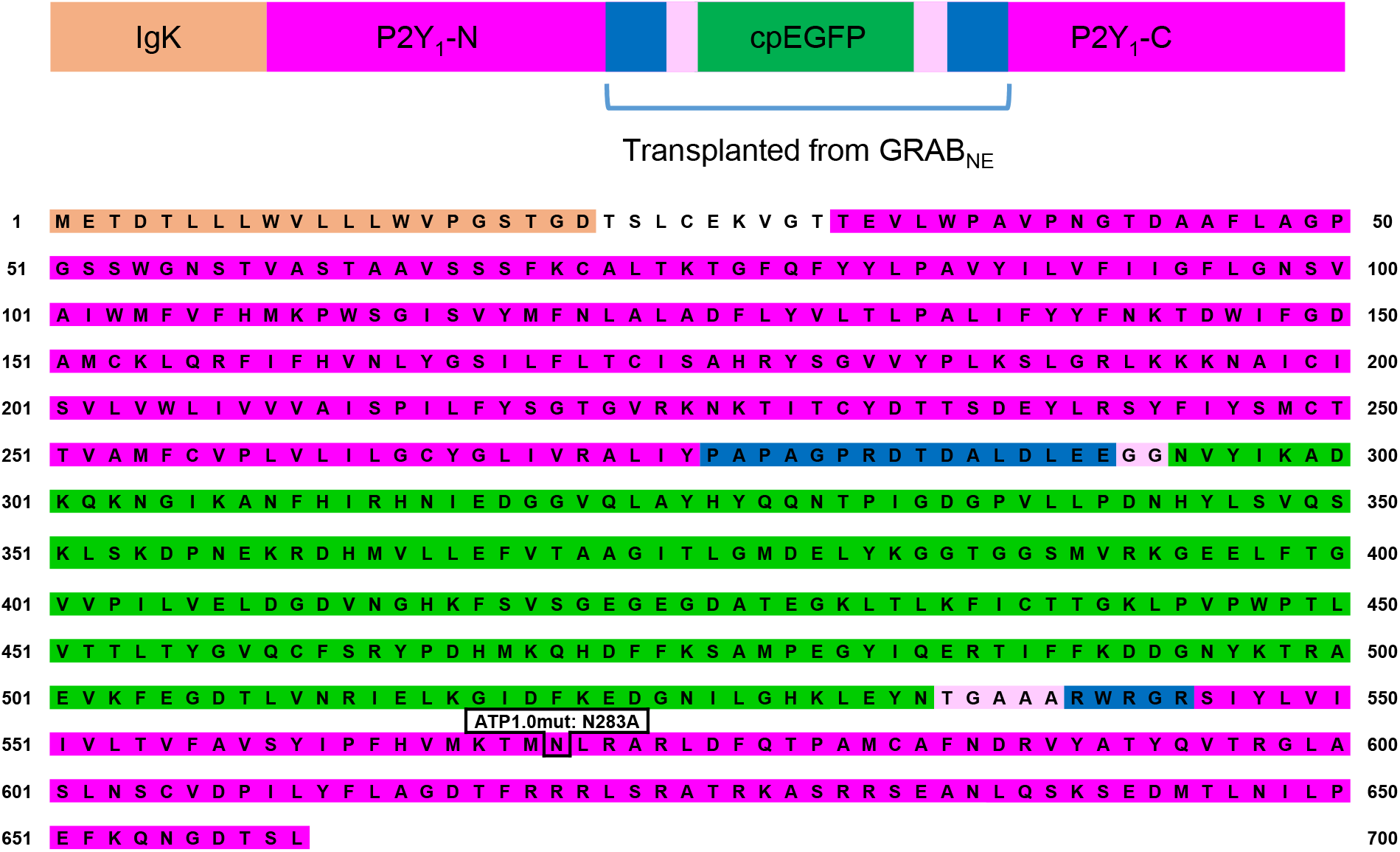
Full amino acid sequence of the ATP1.0 and ATP1.0mut sensors, with the IgK leader sequence, cpEGFP moiety, and N-terminal and C-terminal portions of the P2Y1 receptor indicated.

**Figure S4.**
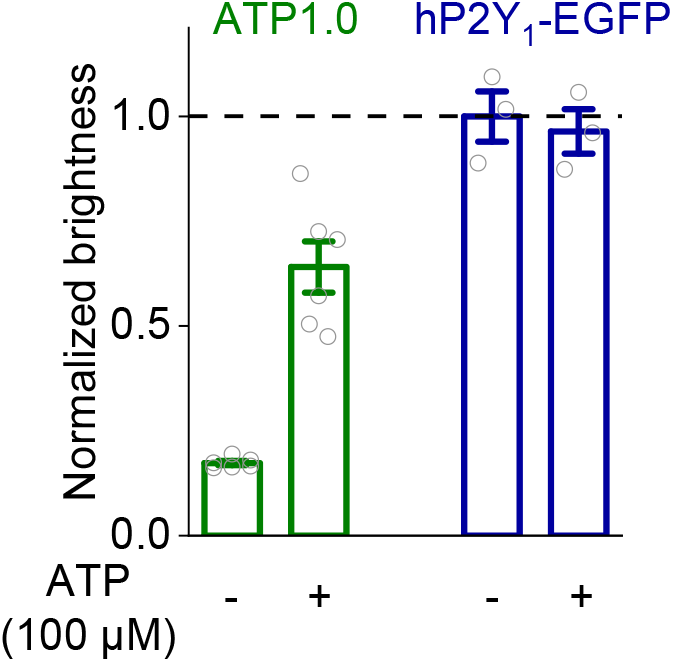
Summary of the normalized brightness of ATP1.0 and hP2Y1-EGFP before (−) and after (+) 100 μM ATP application.

**Figure S5.**
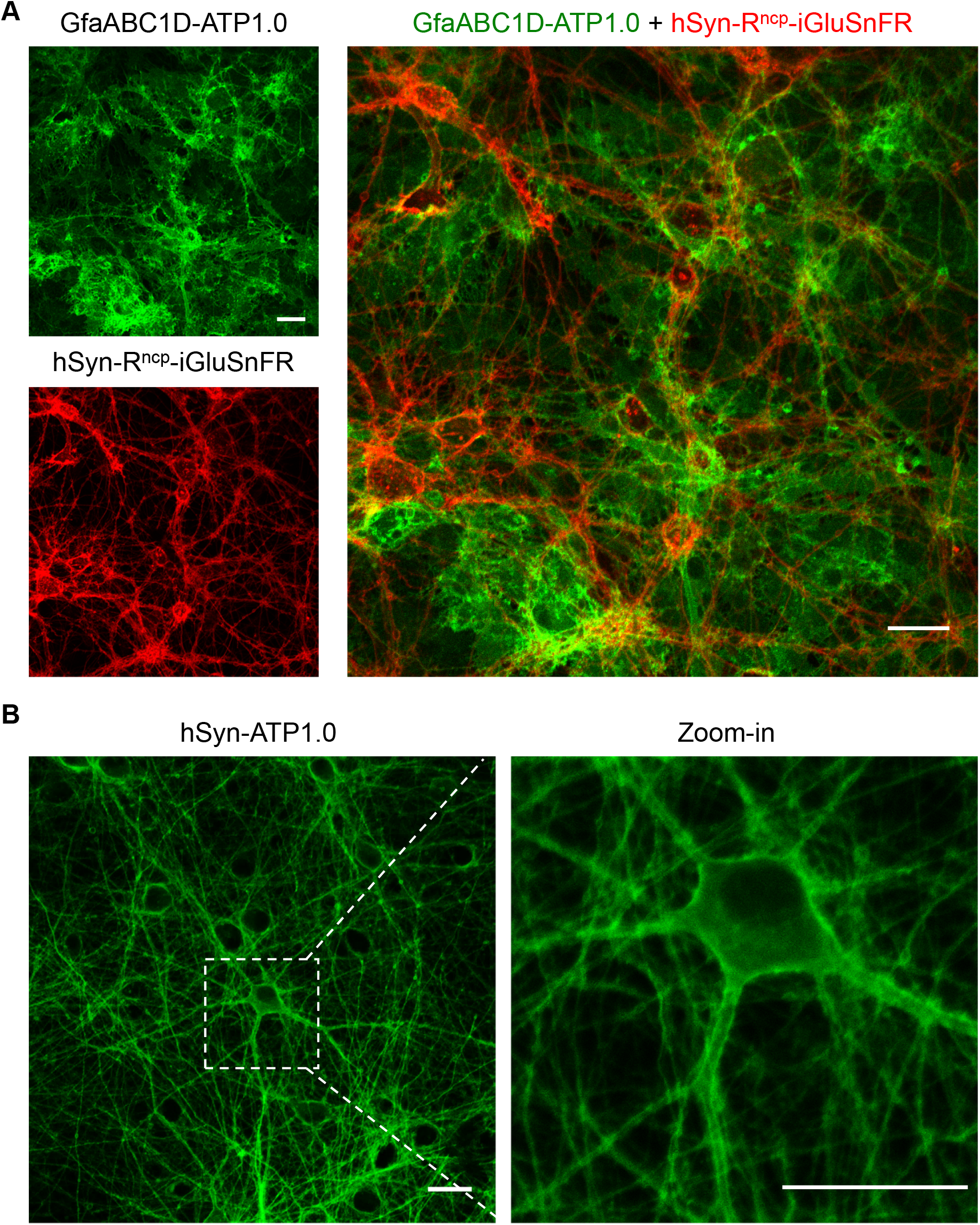
Expression of ATP1.0 in cultured neurons and astrocytes. **(A)** Dual-color imaging of ATP1.0 expressed in astrocytes under the control of the astrocyte-specific GfABC1D promoter and R^ncp^-iGluSnFR expressed in neurons under the control of the hSyn promoter. **(B)** ATP1.0 was expressed in cultured rat cortical neurons under the control of the neuro-specific hSyn promoter. Scale bars represent 30 μm.

**Figure S6.**
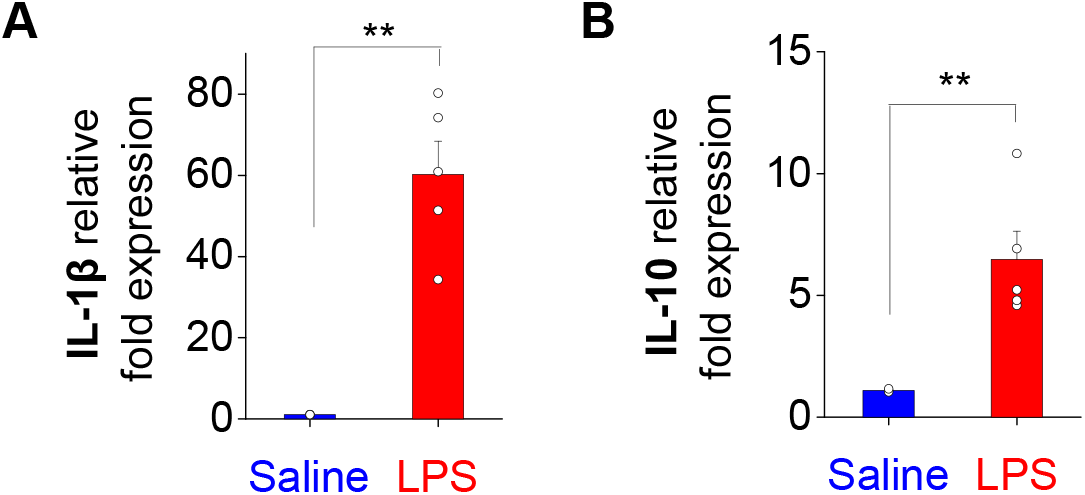
Expression levels of IL-1β and IL-10 in the mouse brain 24 h after an intraperitoneal injection of saline or lipopolysaccharides (LPS) to induce systemic inflammation. Expression of the inflammatory cytokines IL-1β **(A)** and IL-10 **(B)** were measured in the brains of saline- and LPS-injected mice. GAPDH expression was used as an internal control for calculating the fold change in expression. \*\**p*<0.01 (Student’s *t*-test).

## MATERIALS AND METHODS

### Molecular biology

Plasmids were generated using Gibson assembly. DNA fragments were generated using PCR amplification with primers (Tsingke) with ~25-bp overlap, and all sequences were verified using Sanger sequencing. All cDNAs encoding the candidate GRAB_ATP_ sensors were cloned into the pDisplay vector (Invitrogen) with an upstream IgK leader sequence and a downstream IRES-mCherry-CAAX cassette (to label the cell membrane). The cDNAs encoding the ATP receptor subtypes were amplified from the human GPCR cDNA library (hORFeome database 8.1), and the third intracellular loop (ICL3) of each ATP receptor was swapped with the corresponding ICL3 in the GRAB_NE_ sensor. The swapping sites in the P2Y_1_ receptor and the amino acid composition between the P2Y_1_ receptor and the ICL3 of GRAB_NE_ were then screened. The plasmids used to express the GRAB_ATP_ sensors in mammalian neurons and astrocytes were cloned into the pAAV vector under the control of human synapsin promoter (hSyn) or the GfaABC1D promoter, respectively. The plasmids used to express the GRAB_ATP_ sensors in zebrafish were cloned into Elval3: Tet^off^ vectors.

The pm-iATPSnFR1.0 sensor was a gift from Baljit Khakh (Addgene plasmid #102548). The ecAT3.10 sensor was a gift from Mathew Tantama (Addgene plasmid #107215). The SF-iGluSnFR.A184V sensor was a gift from Loren Looger (Addgene plasmid #106175). The R^ncp^-iGluSnFR sensor was a gift from Robert Campbell (Addgene plasmid #107336) and was subcloned into the pAAV-hSyn vector. Finally, the plasmid encoding TeNT was a gift from Dr. Peng Cao and was subcloned into the pAAV-CAG vector.

### Cell cultures, zebrafish, and mice

HEK293T cells and primary neuron-glia co-cultures were prepared and cultured as described previously (Peng et al., 2020). In brief, HEK293T cells were cultured at 37°C in 5% CO_2_ in DMEM (Biological Industries) supplemented with 10% (v/v) fetal bovine serum (FBS, Gibco) and 1% penicillin-streptomycin (Biological Industries). Rat primary neuron-glia co-cultures were prepared from 0-day old (P0) rat pups (male and female, randomly selected) purchased from Charles River Laboratories (Beijing, China). Cortical or hippocampal cells were dissociated from the dissected brains in 0.25% Trypsin-EDTA (Gibco) and plated on 12-mm glass coverslips coated with poly-D-lysine (Sigma-Aldrich) in neurobasal medium (Gibco) containing 2% B-27 supplement (Gibco), 1% GlutaMAX (Gibco), and 1% penicillin-streptomycin (Gibco). Based on glial cell density, after approximately 4 days in culture (DIV4) cytosine β-D-arabinofuranoside (Sigma) was added to the hippocampal cultures in a 50% growth media exchange, with a final concentration of 2 μM.

Rat primary astrocytes were prepared as previously described (Schildge et al., 2013). In brief, the cortex or hippocampi were dissected from P0 rat pups, and the cells were dissociated using trypsin digestion for 10 mins at 37°C and plated on a poly-D-lysine‒coated T25 flask. The plating and culture media contained DMEM supplemented with 10% (v/v) FBS and 1% penicillin-streptomycin. The next day, and every 2 days thereafter, the medium was changed. At DIV 7-8, the flask was shaken on an orbital shaker at 180 rpm for 30 min, and the supernatant containing the microglia was discarded; 10 ml of fresh astrocyte culture medium was then added to the flask, which was shaken at 240 rpm for ≥6 h to remove oligodendrocyte precursor cells. The remaining astrocytes were dissociated with trypsin and plated on 12-mm glass coverslips in 24-well plates containing culture medium. Both the neurons and astrocytes were cultured at 37°C in 5% CO_2_.

For zebrafish experiments, zebrafish larvae at 4-6 days post-fertilization (4-6 dpf) were used for all experiments in this study. As the sex of zebrafish cannot be determined in the larval stage, sex discrimination was not a factor in our study. Wild-type (AB background) and *Tg(coro1a: DsRed)* zebrafish strains were used in this study. Adult zebrafish and larvae were maintained and raised under standard laboratory protocols (Yu et al., 2010), and all procedures were approved by the Institute of Neuroscience, Chinese Academy of Sciences.

Wild-type C57BL/6J mice were housed under a 12-h/12-h light/dark cycle. All protocols for animal surgery and maintenance were approved by the Animal Care and Use Committees at Peking University and the Chinese Institute for Brain Research, and were performed in accordance with the guidelines established by the US National Institutes of Health. Adult mice (>6 weeks of age) were used for the *in vivo* experiments.

### AAV virus preparation

The following AAV viruses were used to infect cultured cells and for *in vivo* expression (all packaged at Vigene Biosciences): AAV2/9-hSyn-ATP1.0, AAV2/9-GfaABC1D-ATP1.0, AAV2/9-hSyn-ATP1.0mut, AAV2/9-GfaABC1D-ATP1.0mut, AAV2/9-CAG-EBFP2-iP2A-TeNT, AAV2/9-SF-iGluSnFR.A184V, and AAV2/9-hSyn-R^ncp^-iGluSnFR.

### Expression of GRAB_ATP_ in cultured cells and *in vivo*

For screening, HEK293T cells expressing the candidate GRAB_ATP_ sensors were plated in 96-well plates (PerkinElmer). For confocal imaging, HEK293T cells were plated on 12-mm glass coverslips in 24-well plates and grown to 60-80% confluence for transfection. Cells were transfected using a mixture containing 1 μg DNA and 1 μg PEI for 4-6 h and imaged 24-48 hours after transfection. For diffuse *in vitro* expression, the viruses were added to neuron-glia co-cultures or cultured astrocytes at DIV 5-9, and the cells were characterized ≥48 hours after infection; DIV ≥13 cells were used for physiological analyses.

For *in vivo* expression in zebrafish, plasmids encoding either ATP1.0 or ATP1.0mut were co-injected (25 ng/μl) with *Tol2* transposase mRNA (25 ng/μl) into one-cell stage wild-type (AB background) or *Tg(coro1a: DsRed)* embryos.

To express GRAB_ATP_ in mice *in vivo*, the mice were anesthetized with an i.p. injection of Avertin (500 mg/kg, Sigma); the skin was retracted from the head, and a metal recording chamber was affixed. After the mice recovered for 1-2 days, the mice were re-anesthetized, the cranial window on the visual cortex was opened, and 400-500 nl of AAV was injected using a microsyringe pump (Nanoliter 2000 injector, WPI) at the following coordinates: AP: −2.2 mm relative to Bregma, ML: 2.0 mm relative to Bregma, and DV: 0.5 mm below the dura at an angle of 30°. A 4 mm × 4 mm square coverslip was used to replace the skull after AAV injection, and *in vivo* two-photon imaging was performed 3 weeks after injection.

### Confocal imaging of cultured cells

Before imaging, the culture medium was replaced with Tyrode’s solution contained (in mM): 150 NaCl, 4 KCl, 2 MgCl_2_, 2 CaCl_2_, 10 HEPES, and 10 glucose (pH 7.3-7.4). For inducing cell swelling, the hypotonic Tyrode’s solution (osmolality: 130 mOsm/kg) contained (in mM): 50 NaCl, 75 KCl, 2 MgCl_2_, 2 CaCl_2_, 10 HEPES, and 10 glucose (pH 7.3-7.4). HEK293T cells grown in 96-well plates were imaged using an Opera Phenix high-content screening system (PerkinElmer) equipped with a 20x/0.4 NA objective, a 40x/0.6 NA objective, a 40x/1.15 NA water-immersion objective, a 488-nm laser, and a 561-nm laser; green fluorescence (GRAB_ATP_ sensors and P2Y_1_R-EGFP) and red fluorescence (mCherry-CAAX) were recorded using a 525/50-nm and 600/30-nm emission filter, respectively. Cells grown on 12-mm coverslips were imaged using a Ti-E A1 confocal microscope (Nikon) equipped with a 10x/0.45 NA objective, a 20x/0.75 NA objective, a 40x/1.35 NA oil-immersion objective, a 405-nm laser, a 488-nm laser, and a 561-nm laser; blue fluorescence (BFP2-TeNT), green fluorescence (GRAB_ATP_ sensors, iATPSnFR1.0, P2Y_1_R-EGFP, and SF-iGluSnFR.A184V), and red fluorescence (mCherry-CAAX and R^ncp^-iGlu) were recorded using a 450/25-nm, 525/50-nm, and 595/50-nm emission filter, respectively.

The following compounds were applied by replacing the Tyrode’s solution (for imaging cells in 96-well plates) or by either bath application or a custom-made perfusion system (for imaging cells cultured on 12-mm coverslips): ATP (Sigma), ADP (Sigma), AMP (Sigma), adenosine (Ado, Sigma), UDP (Sigma), UTP (Sigma), GTP (Sigma), UDP-glucose (Tocris), MRS-2500 (Tocris), Glu (Sigma), GABA (Tocris), Gly (Sigma), DA (Sigma), NE (Tocris), 5-HT (Tocris), HA (Tocris), ACh (Solarbio), and apyrase (Sigma, 15 U/ml apyrase grade VI plus 15 U/ml apyrase grade VII). Between experiments, the recording chamber was cleaned thoroughly using Tyrode’s solution and 75% ethanol. The micropressure application of drugs was controlled using a Pneumatic PicoPump PV800 (World Precision Instruments). Hypotonic solutions were delivered by perfusion. For mechanical stimulation, a glass pipette was placed above the cultured cells. For field stimulation of cultured neurons, parallel platinum electrodes positioned 1 cm apart were controlled using a Grass S88 stimulator (Grass Instruments), and 1-ms pulses were applied at 80 V. Except where indicated otherwise, all experiments were performed at room temperature.

### In vivo confocal imaging of GRABATP in larval zebrafish

GRAB_ATP_ responses induced by local puffing of drugs was performed in zebrafish larvae expressing either *Elval3: Tet^off^-ATP1.0* or *Elval3: Tet^off^-ATP1.0mut*. GRAB_ATP_ responses and microglia movement following laser ablation‒ induced injury were performed by using *Tg(coro1a:DsRed)* zebrafish larvae expressing *Elval3: Tet^off^-ATP1.0*.

*In vivo* confocal imaging experiments were performed using an FN1 confocal microscope (Nikon) equipped with a 40x (NA 0.8) or 25 x (NA 1.1) water-immersion objective. Before imaging, the larvae were immobilized in 1.2% low-melting point agarose. Time-series imaging was carried out at 28°C using a heating system. A 488-nm or 561-nm excitation laser and a 525/50-nm or 595/50-nm emission filter were used to excite and collect the GFP and DsRed signals, respectively.

To monitor the GRAB_ATP_ responses to locally puffed drugs, the larvae were paralyzed with 1 mg/ml α-bungarotoxin (Tocris), the agorae around the tectum region were removed, and a small incision in the skin around the top tectum was made for introducing the micropipette. The larvae were incubated in external solution (ES) either with or without 90 μM MRS-2500 (Tocris). Local puff application of ES with or without 5 mM ATP (Sigma) was performed using a micropipette with a tip diameter of 1-2 μm introduced via the contralateral optic tectum to the target tectum region. For each zebrafish larva, the solution contained in the micropipette was puffed using 2 pulses of gas pressure (3 psi, 50-ms duration, 1-s interval), with 5 local puffing sessions in total applied at a 2-min interval. Single optical section confocal imaging was performed with an interval of 2.2 s.

To monitor the GRAB_ATP_ responses and microglial dynamics following laser ablation in larval zebrafish, the larvae were paralyzed and imaged as described above. Time-series images were captured before and immediately after laser ablation. For laser ablation, target regions (5 μm in diameter) were illuminated at 800 nm for 7 s using a two-photon laser.

### Two-photon *in vivo* imaging in mice

Two-photon imaging was performed using a FluoView FVMPE-RS microscope (Olympus) equipped a laser (Spectra-Physics). For experiments involving lipopolysaccharides (LPS), 10 mg/kg LPS from *Escherichia coli* O111:B4 (Sigma, L4130) was dissolved in sterile saline and injected intraperitoneally (i.p.) into the mice. ATP1.0 was imaged using a 920-nm laser, and the imaging frequency was set at 32 Hz with 512×512-pixel resolution.

### Data analysis

Imaging data obtained from cultured cells and zebrafish were first processed using ImageJ software (NIH) or MATLAB 2018 (MathWorks); traces were generated using OriginPro 2019 or MATLAB, and pseudocolor images were generated using ImageJ. For the mouse 2-photon microscopy images, 10 images were first averaged and processed using AQuA software in MATLAB, and the detail information regarding individual ATP-release events were plotted using either MATLAB with custom-written programs or OriginPro 2019.

Except where indicated otherwise, all summary data are presented as the mean ± SEM. Groups were analyzed using either the Student’s *t*-test or a one-way ANOVA.

### Data and software availability

The plasmids for expressing ATP1.0 and ATP1.0mut used in this study have been deposited at Addgene (https://www.addgene.org/Yulong_Li/).

